# High-throughput quantitative detection of basal autophagy and autophagic flux using image cytometry

**DOI:** 10.1101/590513

**Authors:** Gopika SenthilKumar, Justin H. Skiba, Randall J. Kimple

**Affiliations:** Department of Human Oncology and UW Carbone Cancer Center – University of Wisconsin School of Medicine and Public Health

**Author notes:** Corresponding author; **Contact information:** Randall J Kimple, Department of Human Oncology, University of Wisconsin School of Medicine and Public Health, University of Wisconsin – Madison, Madison, WI 53705 USA, @kimplerandall. **Author contributions:** GS and JHS performed the developed the project, performed the work, and drafted the manuscript. RJK oversaw all aspects of the work including project definition, methods and analysis, and manuscript writing.

**Keywords:** image cytometer, autophagy, acridine orange

## Abstract

Quantitative assessment of changes in macro-autophagy is often performed through manual quantification of the number of LC3B foci in immunofluorescence microscopy images. This method is highly laborious, subject to image-field selection and foci-counting bias, and is not sensitive for analyzing changes in basal autophagy. Alternative methods such as flow cytometry and transmission electron microscopy require highly specialized, expensive instruments and time-consuming sample preparation. Immunoblots are prone to exposure-related variations and noise that prevent accurate quantification. We report a high-throughput, inexpensive, reliable, and objective method for studying basal level and flux changes in late-stage autophagy using image cytometry and acridine orange staining.

**Methods summary:** A high-throughput, inexpensive, reliable, and objective method for studying both basal autophagy and autophagic flux is reported. This approach uses acridine orange staining of late-stage autophagy and image cytometry to quantify autophagy.

---

Macro-autophagy (referred to as autophagy hereafter) is a cellular recycling mechanism used for maintaining cytoplasmic homeostasis and surviving a variety of cellular stresses (1, 2). The most common inducer of autophagy is nutrient deprivation, although pharmacologic activation using the mTOR inhibitor, PP242, also induces autophagy (3) (Figure 1).

**Figure 1.**
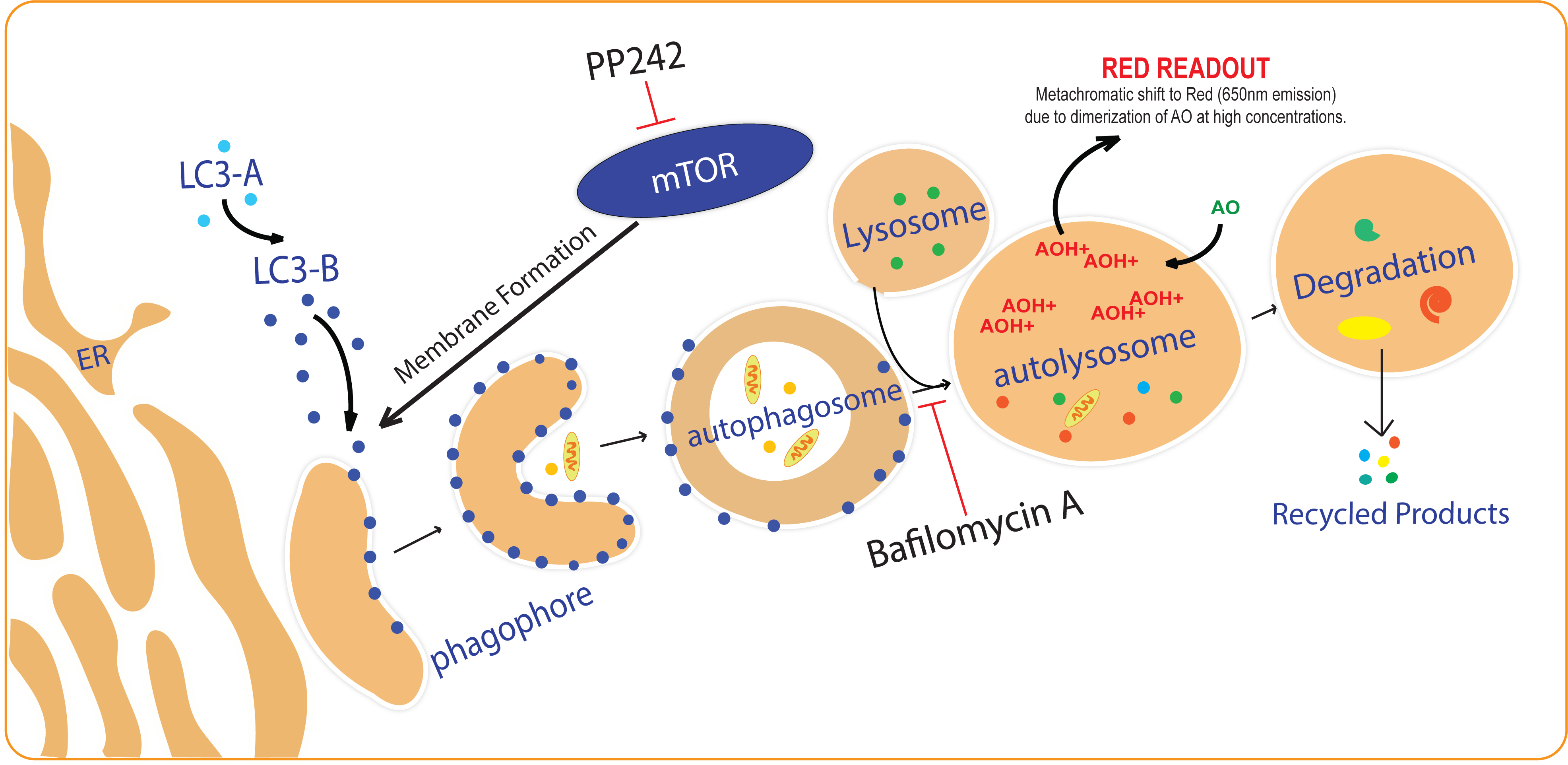
Autophagic Pathway.

The first step in autophagy induction involves the formation of a double membraned autophagosome that engulfs cellular materials and subsequently fuses with the lysosome to form an autolysosome, where cellular materials are degraded (4). Protein light-chain 3B-II (LC3B-II) is a key marker found on autophagosome membranes and its levels are studied using immunofluorescence (IF), immunoblot, flow cytometry, and transmission electron microscopy (TEM) to understand changes in autophagy (2). LC3B-II is degraded upon autolysosome formation, and thus inhibitors of autolysosome formation such as Bafilomycin A1 are used to accumulate LC3-IIB for a standard amount of time to study autophagic flux (2, 5). However, the quantification of immunofluorescence is subject to imaging field-selection bias, counting bias, and observer variations even when using semi-automated quantification methods (6). Immunoblot quantification is subject to incorrect results from minor loading errors and from over-saturation of house-keeping or target proteins in an attempt to distinguish differences in the protein of interest (7). TEM requires specialized equipment and specific expertise for interpreting data and thus can be prone to misinterpretation (8). Flow cytometry requires specialized equipment and time-consuming sample preparation and is better suited for discerning the proportion of a sample that is undergoing autophagy. These methods are summarized and compared in Table 1.

**Table 1.** Comparison of commonly used methods for studying autophagy.

| Analysis Method | Result provided | Sensitivity for Basal Autophagy | Sensitivity for autophagic flux | Assay Time | Sample Prep Cost | Instrument Cost | Quantification Reliability |
| --- | --- | --- | --- | --- | --- | --- | --- |
| Immunofluorescence microscopy | Area or foci of LC3B per cell | + | +++ | ~8hrs | ++ | +++ | ++ |
| Western Blot | Specimen average LC3B expression | + | +++ | ~6hrs | ++ | + | + |
| Flow Cytometry | Average intensity of LC3B or acridine orange per cell | +++ | +++ | ~4+ hrs | ++ | +++ | ++++ |
| Transmission Electron Microscopy | Morphological structures of autophagosomes and autolysosomes in cells | ++++ | ++++ | ~10+ hrs | ++++ | ++++ | ++ |
| Image Cytometer | Average intensity of acridine orange per cell | +++ | +++ | ~30-45mins | + | ++ | ++++ |

Increases in autophagosomes can be caused by either autophagy induction or inhibition of downstream processing such as lysosomal fusion (2, 9) (Figure 1). Therefore, it is important to analyze late stage autophagy, such as changes in autolysosome levels, to directly study autophagic induction. Acridine orange (AO) is a fluorophore that accumulates in a protonated-form inside acidic vesicular organelles (AVO) such as autolysosomes. **At high concentrations, AO dimerizes causing a metachromatic shift from green to red (Figure 1), which can be measured for studying late-stage autophagy. Thomé et al. can be referenced for detailed chemical and concentration changes (10)**. AO can be assessed using fluorescence microscopy or flow cytometry, but these approaches suffer from the limitations described above (Table 1). We have used image cytometry as a quantitative, high-throughput, objective, and inexpensive method (referred to as ‘AO assay’ from hereafter) that can be used to measure changes in basal autophagy and autophagic flux using acridine orange. **The AO assay is a useful screening tool and reliable, unbiased complement to the more qualitative and expensive approaches described above for studying autophagy**.

UM-SCC47 cells were cultured in black-wall, clear-bottom 96-well plates (the outer wells were left empty as they were prone to noise/ incorrect read-outs) and on 8-well IF slides. IF was used as a qualitative, established comparison for the AO assay. Cells were treated with DMSO, PP242 (500nM or 2μM) both with and without Bafilomycin A1 (Baf, 100nM) for 24hours. For IF, 24 hours post-treatment, slides were fixed and tagged with LC3B (D11) XP Rabbit mAb (Cell Signaling Technology #3868), fluorescently conjugated (Alexa Fluor 488), stained with DAPI (Invitrogen Fluoromount-G with DAPI), and 3 random fields imaged using a 40x objective (Olympus BX41 inverted microscope with XM10 camera). The area of LC3B foci was quantified using **FIJI** and normalized to cell count (11). A 2-way ANOVA multi-variance test (alpha .05) was performed using GraphPad Prism (GraphPad Software, Inc., La Jolla, CA). Both basal autophagy (Baf-) and autophagic flux (Baf+ minus Baf-) was assessed (Figure 2A/B). A different IF slide was washed with 1X phosphate buffered saline at pH 7.4 (1xPBS) and stained with 1μg/ml acridine orange solution diluted in 1xPBS. The slide was cover-slipped and immediately imaged as described above using a 20x objective (Figure 2C/D).

**Figure 2.**
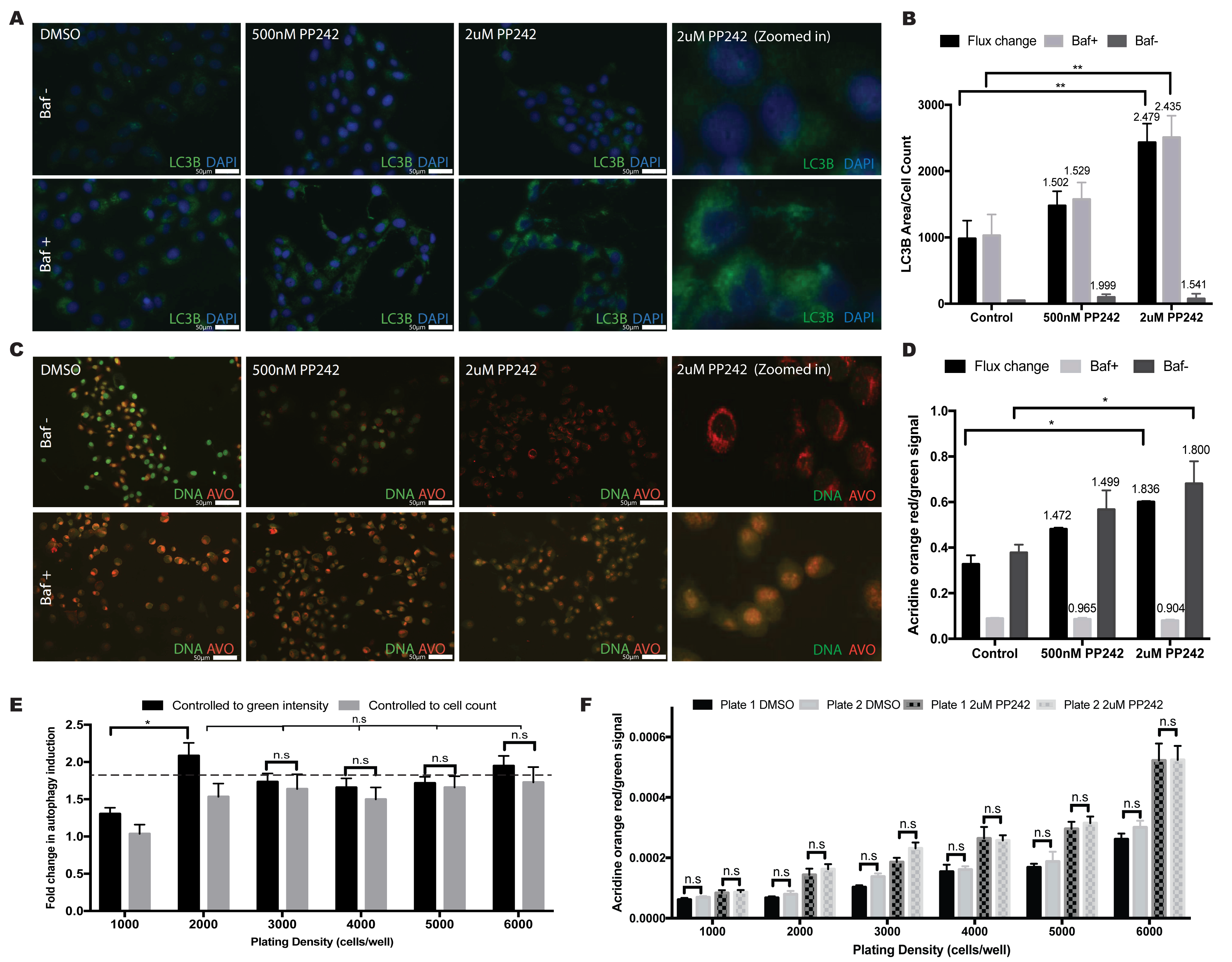
The analysis of acridine orange signal using the image cytometer provides an objective, high-throughput method for analyzing basal and flux changes in autophagy. (A) Representative 40x magnification IF images tagged with LC3B antibody following treatments as indicated. (B) Average area of LC3B foci per cell [+/ − 95% confidence interval] measured using **FIJI**. The values above the bars indicate fold change. (C) Representative 40x images of AO stained IF slides following treatments as indicated. (D) Average AO red/green signal ratio [+/ − 95% confidence interval] measured using image cytometer. The values above the bars indicate fold change. (E) Fold change results following 2μM PP242 treatment are consistent regardless of the normalization method used (cell count vs. green intensity) at optimal plating densities.**Dashed line shows the expected fold change from figure 2D**. (F) There are no significant differences in the average red/green signal ratio following 2μM PP242 treatment within the same treatment groups across multiple plates at varying cell densities.

For the image-cytometer based AO Assay, 24 hours post-treatment, 96-well plates were washed three times (100μL/well) with 1xPBS. 50μL of 1μg/ml AO was added to each well and the plates were incubated at room temperature for 30 minutes. Cells were then washed two times (100μL/well) with 1X PBS. 50μL of 1X PBS was added per well, and then the plates were read on the SpectraMax i3 Multi-Mode Microplate Reader Platform with MiniMax 300 Imaging Cytometer (Molecular Devices, Sunnyvale, CA) using the SoftMax Pro software (v6.4).

The monochromator and MiniMax settings were used on the software (Supplemental Figure 1). The monochromator setting was used to measure excitation/emission wavelengths of 500/526 (green) **to assess intensity of unprotonated, diffuse AO staining DNA (non-autophagic staining), and 460/650 (red) to assess intensity of dimerized, protonated AO concentrated in AVOs (autophagic staining)** (10). The MiniMax setting was used to count the number of cells per well as we have previously described (12). The red intensity signal per well was divided by the green intensity or the cell count to assess the level of autophagy and to assess the efficacy of the two normalization methods: controlling to green intensity as recommended by Thome et al. vs. controlling to cell count as previously done by Fowler et al. (10, 12). The statistical analysis performed on the IF data was performed on the AO assay recorded values. Basal level autophagy and autophagic flux (Baf-minus Baf+) was graphed along with a 95% confidence interval.

As shown in Figure 2A/B, imaging LC3B demonstrates a dose-dependent increase in autophagic flux but does not demonstrate a dose response in basal autophagy. However, the quantified results should be viewed with caution as slight changes in the threshold used for quantification can drastically change the results (Supplemental Figure 2). We recommend using the IF images as a qualitative comparison. The graph in Figure 2B shows a significant increase in autophagic flux, but not basal level autophagy, following 2μM PP242 treatment. The higher fold change in basal autophagy after 500nM PP242 treatment compared to 2μM treatment as well as the low LC3B foci areas reflect the insensitivity of the assay to analyzing basal level changes.

Fluorescent images of AO stained cells show a dose-dependent increase in basal autophagy (Figure 2C, top row) as illustrated by increased red intensity and decreased green intensity. In the Baf-group, the increase in red cytoplasmic intensity in conjunction with a decrease in green nuclear intensity with increasing doses of PP242 reflects a PP242 dose-dependent increase in basal autophagy (10). In the Baf+ group, no significant differences are observed between groups since Baf inhibits late stage autophagy (i.e. accumulation of AVOs). Regardless, the Baf can be used for quantitatively studying autophagic flux (red/green ratio of Baf+ subtracted from Baf-, Figure 2D). The graph shows basal autophagy and autophagic flux measured by the image cytometer using AO (plating density: 5000 cells per well). An increase in both flux change and basal autophagy is seen following 2μM PP242 treatment, but not after 500nM PP242 treatment. This data is consistent with the images in Figure 2C where a decrease in green but only a slight increase in red staining is seen after 500nM treatment. However, in the 2μM group, a clear shift from green to red is observed. The same trend is seen in Figure 2A and 2B as well, with a clear increase in LC3B puncta following 2μM PP242 treatment, but not much change after 500nM treatment (Baf+ group). **A correlation of autophagic flux between LC3B and AO shows an R**^**2**^ **value of** .**9508 (Supplemental Figure 3) thereby confirming the sensitivity and accuracy of the AO assay. The efficacy of this assay was also confirmed using other cancer cell lines and other autophagy inducers/inhibitors (Data not shown)**.

To assess the effect of cell plating density and compare different normalization methods, we repeated the experiment using a range of plated cell densities (Figure 2E). At low densities, inconsistent results are seen (1,000 and 2,000 cells/well) while better consistency in this cell line is seen at plating densities between 3,000-6,000 cells/well. Similar results were seen independent of the normalization method used: green nuclear intensity vs. cell count. Due to the significantly faster analysis using green intensity, this is our preferred approach. We also compared results across different plates and saw no consistent differences (Figure 2F).

This data demonstrates the value of image cytometry of AO to assess changes in basal autophagy and autophagic flux. The inexpensive nature of this assay, its ability to be used in high-throughput screens, and the objective nature of the data analysis make this assay a useful tool and complement to more qualitative approaches to studying autophagy.

## Notes

**Conflict of interest:** The authors report no financial or non-financial relationships exist that potentially conflict with the subject matter or materials.

**Disclosure:** Supported in part by grants from the American Cancer Society, the University of Wisconsin Carbone Cancer Center Support Grant (P30 CA014520), and the Wisconsin Head and Neck Cancer SPORE Grant (NIH P50 DE026787).

#### Summary of Updates

Revision in response to peer-review.

