## Supplementary figures and images for "High-throughput quantitative detection of basal autophagy and autophagic flux using image cytometry"

### Supplemental Figure 1

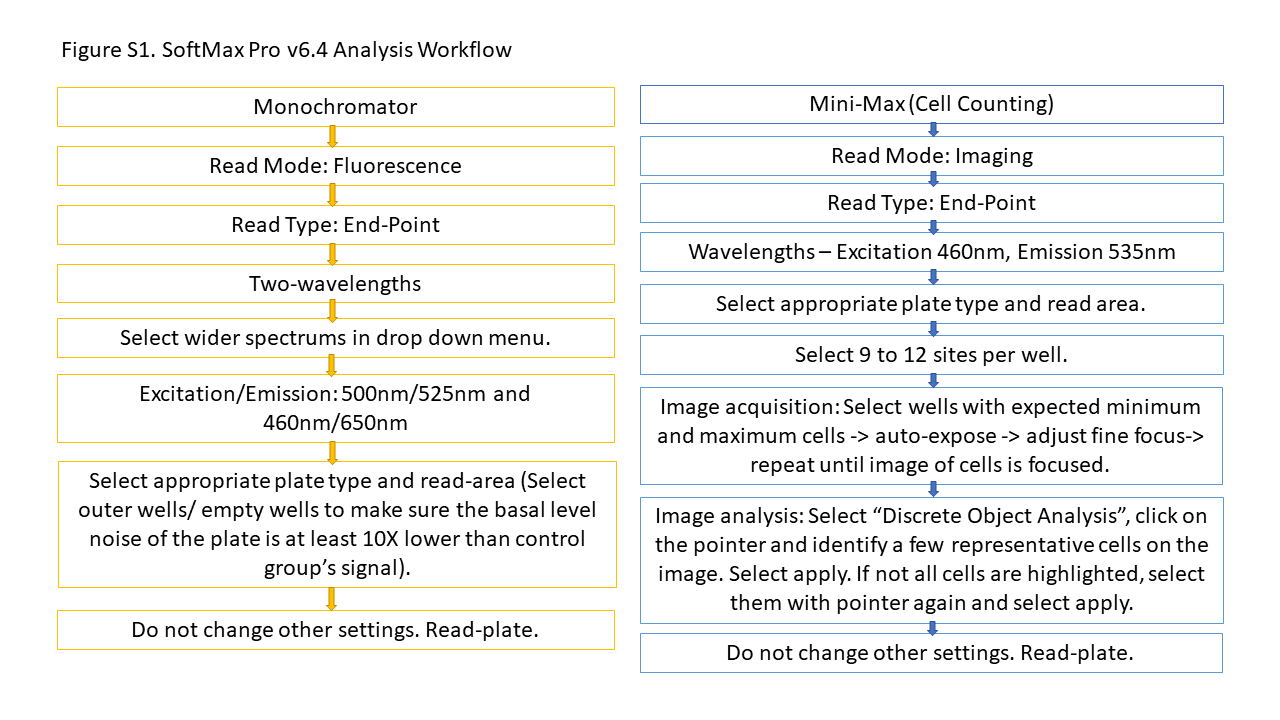

### Supplemental Figure 2

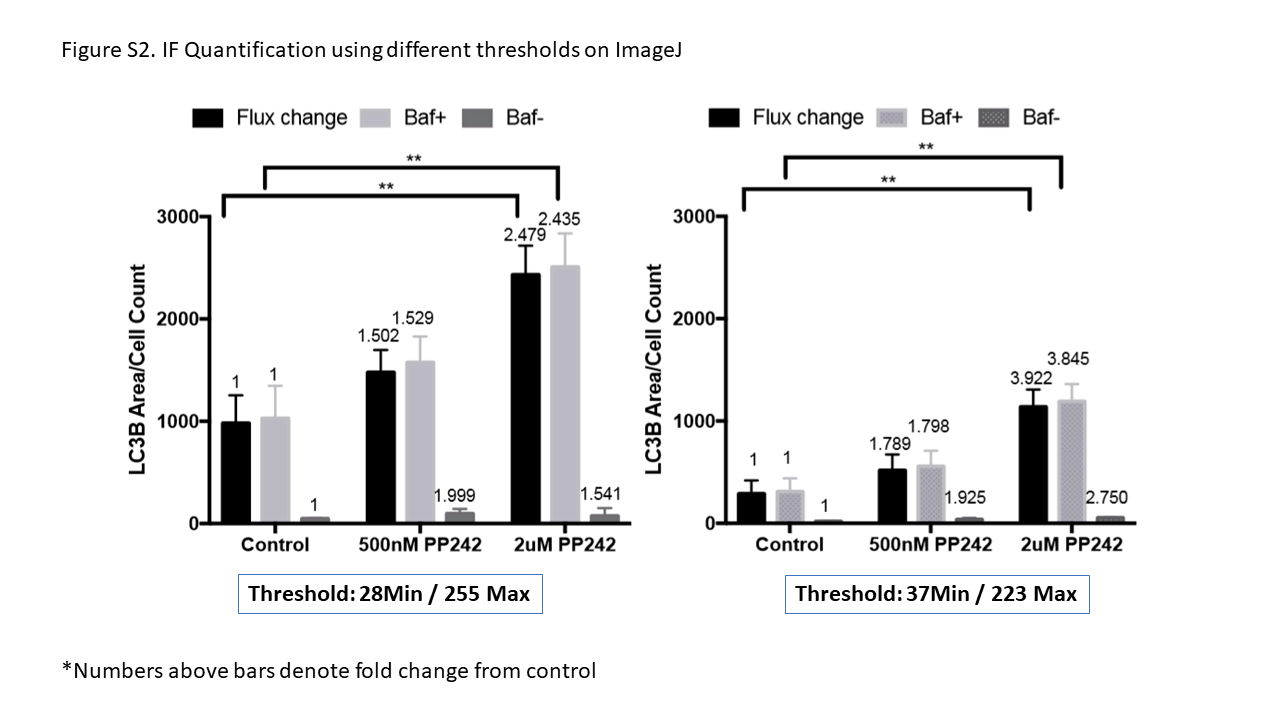

### Supplemental Figure 3

Figure S3. Correlation between IF flux change and AO assay flux change

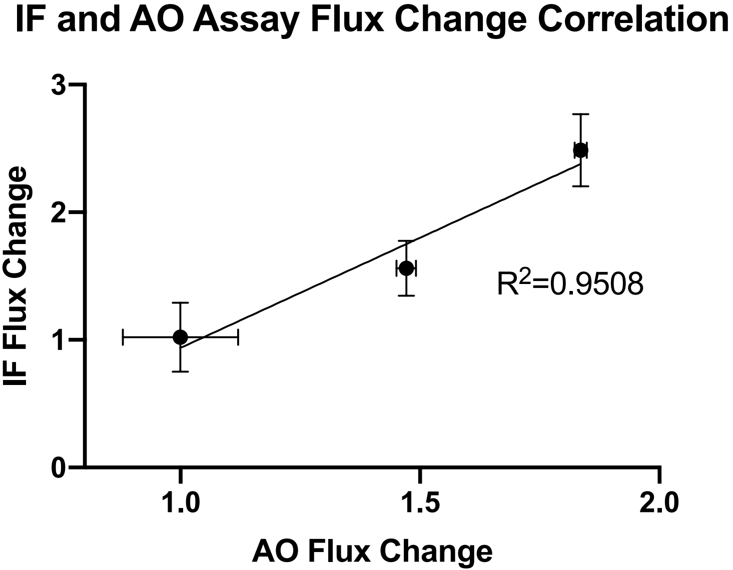
